# Endoplasmic Reticulum Associated Degradation (ERAD) Deficiency Promotes Mitochondrial Dysfunction and Transcriptional Rewiring in Human Hepatic Cells

**DOI:** 10.1101/2020.04.22.053546

**Authors:** Xiaoqing Yang, Qingqing Liu, Guangyu Long, Yabin Hu, Zhenglong Gu, Yves R. Boisclair, Qiaoming Long

## Abstract

Mitochondrial dysfunction has been associated with a variety of human diseases including neurodegeneration, diabetes, non-alcohol fatty liver disease (NAFLD) and cancer, but its underlying causes are incompletely understood. Endoplasmic reticular associated degradation (ERAD) is a protein quality control process essential for maintaining ER homeostasis. Using the human hepatic cell line HepG2 as a model, we show here that ERAD is critically required for mitochondrial function in mammalian cells. Pharmacological inhibition or genetic ablation of ERAD increases cell death under both basal conditions and in response to proinflammatory cytokines. Decreased viability of ERAD-deficient HepG2 cells was traced to impaired mitochondrial functions including reduced ATP production, enhanced reactive oxygen species (ROS) accumulation and increased mitochondrial outer membrane permeability (MOMP). Transcriptome profiling reveals widespread down-regulation in the expression of genes underpinning mitochondrial functions, and up-regulation in the genes with association to tumor growth and aggression. These results highlight a critical role for ERAD in maintaining mitochondrial functional and structural integrity and raise the possibility to improve cellular and organismal mitochondrial function via enhancing cellular ERAD capacity.

## Introduction

The endoplasmic reticulum (ER) serves several critical cellular functions, including Ca2+ storage, lipid synthesis and the folding of protein molecules destined for the secretory pathway. Impaired ER function or ER stress has been implicated in a increasing number of chronic human diseases, including obesity, diabetes, neurodegeneration, non-alcohol fatty liver disease (NAFLD) and cancer [1-3]. ER stress is an abnormal homeotic condition characterized by the accumulation of unfolded or misfolded proteins in the ER lumen [4]. The classical view is that ER stress activates the unfolded protein response (UPR), which is coordinated by the ER stress sensors, PERK, IRE1a and ATF6 [5]. Activation of these sensors reduces protein translation and enhances protein folding and ER-associated degradation (ERAD). Failure of these actions to restore homeostasis results in sustained ER stress, activation of the CCAAT-enhancer-binding protein homologous protein (CHOP) pathway and apoptosis [6].

The ER is a well-characterized ‘functional partner’ of many organelles and dysregulation of these interactions may impair cellular function [7]. The interactions between the ER and mitochondria, an organelle essential for life and death of eukaryotic cells [8-10], provide a good example of the importance of this partnership. The ER and mitochondria form tight physical contacts at the mitochondria associated membranes (MAMs) to regulate several key cellular processes, including Ca2+ homeostasis, lipid exchange, steroid biosynthesis and apoptosis [11]. The MAMs were also found to function as a signaling hub for mitochondria fission as well as autophagy, key events involved in mitochondrial quality control [12]. Over the last few years, biochemical and genetic approaches have provided evidence how these two organelles interact physically and functionally to coordinate various cellular functions [13, 14]. However, less is known about the potential pathophysiological consequences of an impaired UPR pathway in the ER on mitochondria function and vice versa.

We have recently reported that pharmacological and genetic disruption of ERAD, an ER protein quality control process, impairs glucose stimulated insulin secretion in cultured rat pancreatic Β-cell line and in mice [15]. By utilizing an ERAD-deficient hepatic cell line model, we demonstrate here that ERAD-deficiency causes structural and functional damages to mitochondria. Consequently, ERAD-deficient cells show impaired ATP production, increased mitochondrial outer membrane permeability (MOMP) and enhanced sensitivity to tumor necrosis factor alpha (TNFα)-induced cell death. These consequences of ERAD deficiency were associated with an altered gene expression profile characterized by the selective down-regulation of mitochondria-related and up- regulation of tumor growth and aggression-related genes. These results indicate that ERAD is critically required for maintaining normal mitochondrial function and ERAD- deficiency may potential mammalian cells to adopt a tumorigenic molecular signature by activating the mitochondrial retrograde signaling pathway.

## Materials and Methods

### Cell culture, plasmid transfection, fluorescent imaging and CRISPR/Cas9 mediated gene editing

HepG2 cells were cultured in DMEM medium supplemented with 15% FBS,100U/ml penicilin,100 μg/ml streptomycin at 37°C in an humidified CO_2_ incubator. For visualization of mitochondria or monitoring ER stress, the expression plasmids p-mtGFP containing mitochondria-targeted green fluorescent protein (GFP), or pCAX-F-XBPDBD- venus [16] was transfected into HepG2 cells using Lipofectamine 3000. GFP images were acquired with a FluoviewFV1000 Olympus confocal microscope analyzed by the software Image J (NIH Image).

For Cas9-mediated gene editing, sgRNAs flanking exon 6 were designed using the Optimized CRISPR Design tool (http://crispr.mit.edu/) and cloned into the expression vector pGL3-U6-2sgRNA. To generate SEL1L-deficient HepG2 cell lines, the sgRNA expression vector and the Cas9-expression plasmid pST1374-NLS-3xflag-linker-Cas9 [17] were co-transfected into HepG2 cells using Lipofectamine 3000. After transfection, puromycin (5 μg/ml) was added to the culture medium to select for puromycin-resistant cells. A week later, puromycin-resistant HepG2 colonies were picked, expanded and analyzed by PCR using primers F (5’-GGCAATGATTGACTCAAGTTGTA) and R1 (5’- GCCTTTTGGAGATACCGATATG) flanking exon 6 and exon 6-specific primer R2 (5’- GGCAGAGACTGGAGTGCTCACT).

### Cell death and viability assay

Wild type and Sel1L-deficient HepG2 cells were seeded into a 6-well plate at an initial density of 1×10^5^/well and allowed to recover for 12 hours. Propidium iodide (PI) and annexin V were then added into the culture medium (10 ng/ml) and cell death was monitored in real-time over the next 72 hours using an Incucyte Zoom system. Cell viability was assessed by MTT assay or by fluorescence activated cell sorting (FACS) using a Beckman Counter Flow Cytometer.

### ROS, cytosolic and mitochondrial Ca^2+^ measurement

HepG2 cells were seeded into 4-well glass-bottomed chambers (Nunc,IL) at a density of 1×10^5^/well in DMEM medium and cultured for 24 h. Cytosolic and mitochondrial Ca^2+^and intracellular ROS were measured by adding the freshly prepared fluorescent dye Fluo4/AM, Rhod2 AM (Invitrogen) and Mitosox into the culture medium followed by cell imaging with a FluoviewFV1000 Olympus confocal microscope.

### Mitochondrial transmembrane potential (MTP), oxidative phosphorylation (OXPHOS) and glycolytic functional analysis

MTP was determined using a JC-1 mitochondrial membrane potential assay kit (Abcam). Oxidative phosphorylation and glycolysis were determined using a Seahorse XFe24 extracellular flux analyzer (Seahorse Bioscience) according to the manufacturer’s protocols and as previously described [18]. Briefly, a total of 4 x10^4^ cells were seeded in each well of Seahorse XF24 cell culture microplate and cultured for 16 hours in complete medium. The oxygen consumption rate (OCR) and extracellular acidification rate (ECAR) were assayed using the XF Cell Mito Stress Test Kit and XF Glycolysis Stress Test Kit, respectively, coupled with the XF24-3 Extracellular Flux Assay Kit. All OCR and ECAR measurements were normalized to protein content.

### Western blotting and probing

Protein isolation, gel electrophoresis and blotting were performed essentially as described [15]. After blocking with 5% skim milk, membranes were incubated at 4°C overnight with primary antibodies aginst: p-JNK (Santa Cruz, 1:800), Caspase-1 (Santa Cruz, 1:800), JNK1 (Santa Cruz, 1:800), RIPK1 (Affinity, 1:1000), Cytc (CST, 1:1000), MCU (Sigma, 1:800) and β-actin (Sigma, 1:1000). Signals were developed by incubating with 1/5000 fold dilution of anti-rabbit or anti-mouse secondary antibodies for 1 h at room temperature and quantified using the Image Lab Software on a ChemiDoc XRS system (Bio-Rad).

#### RNA-Seq and Bioinformatics analyses

Total RNA was isolated from HepG2 cells grown under standard conditions using Trizol Reagent (Invitrogen). RNA-Seq was performed by GENEWIZ Co., Ltd. (Suzhou, China). In brief, libraries were prepared using the VAHTS mRNA-seq V2 Library Prep Kit for Illumina, according to the manufacturers’ instructions. Sequencing libraries were run on an Illumina Novaseq platform with the 2 × 150-bp paired-end protocol. These RNA-Seq data has been deposited onto public repository of Genome Sequence Archive (GSA) database in BIG Data Center (BioProject: PRJCA002400; GSA accession number: CRA002460).

Raw reads yielded by paired-end transcriptome sequencing were mapped to the human reference genome (GRCh37/hg19) by tophat2 (version 2.1.1) [19]. Read counts were obtained from BAM files using samtools (version 1.9) {Li, 2009 #85} and gfold (version 1.1.4) [20], and then converted into log2-counts-per-million (logCPM) values to quantify the gene expression level by the R package edgeR (version 3.26.8) [21]. Statistical significances of differentially expressed genes were determined by the empirical Bayes method powered by the R package limma (version 3.40.6) [22] with *p* < 0.005 as the statistical cutoff. Programs written in R language (version 3.6.1) use library extension to pull in functions from non-core packages from bioconductor repository (https://www.bioconductor.org/).

ToppGene Suite (https://toppgene.cchmc.org/) [23] was used to perform the enrichment analysis of differentially expressed genes for Gene Ontology (GO) biological process (Gene_Ontology_Consortium, 2015), Reactome pathway [24], and Kyoto Encyclopedia of Genes and Genomes (KEGG) pathway [25]. Gene set enrichment analysis (GSEA, version 4.0.3) [26] was performed to identify the priori defined gene set associated with transcriptome differences between phenotype (*SEL1L-*deficient vs intact wild type). The c6.all.v7.0.symbols.gmt (oncogenic signatures) and c2.cgp.v7.0.symbols.gmt (chemical and genetic perturbations) gene set libraries were used as the reference gene set collection for enrichment analysis. The statistical cutoff for this analysis was set at *P* < 0.05.

### Statistical analyses

Data were analyzed by Student’s t-test or two-way repeated ANOVA using Graphpad (7.0) with statistical significance set at p<0.05. Data are present as mean ± standard error.

## Results

### Pharmacological inhibition of ERAD perturbs HepG2 cell mitochondrial function and reduces cell viability

We recently showed that ERAD disruption reduces ATP production in rat pancreatic Β- cell line INS-1, indicating that ERAD deficiency impairs mitochondrial respiratory function [15]. To determine whether this phenomena extends to other mammalian cells, we treated human hepatic HepG2 cells with the ERAD inhibitor EerI. EerI treatment caused dose- and time-dependent decreases in ATP levels (Fig. 1A-B) and cell viability (Fig. 1C-D). These dose and time-dependent EerI effects were associated with reciprocal increases in mitochondrial levels of calcium (Ca2^+^) (Fig. 1E-F) and ROS (Fig. 1G-H).

**Figure 1.**
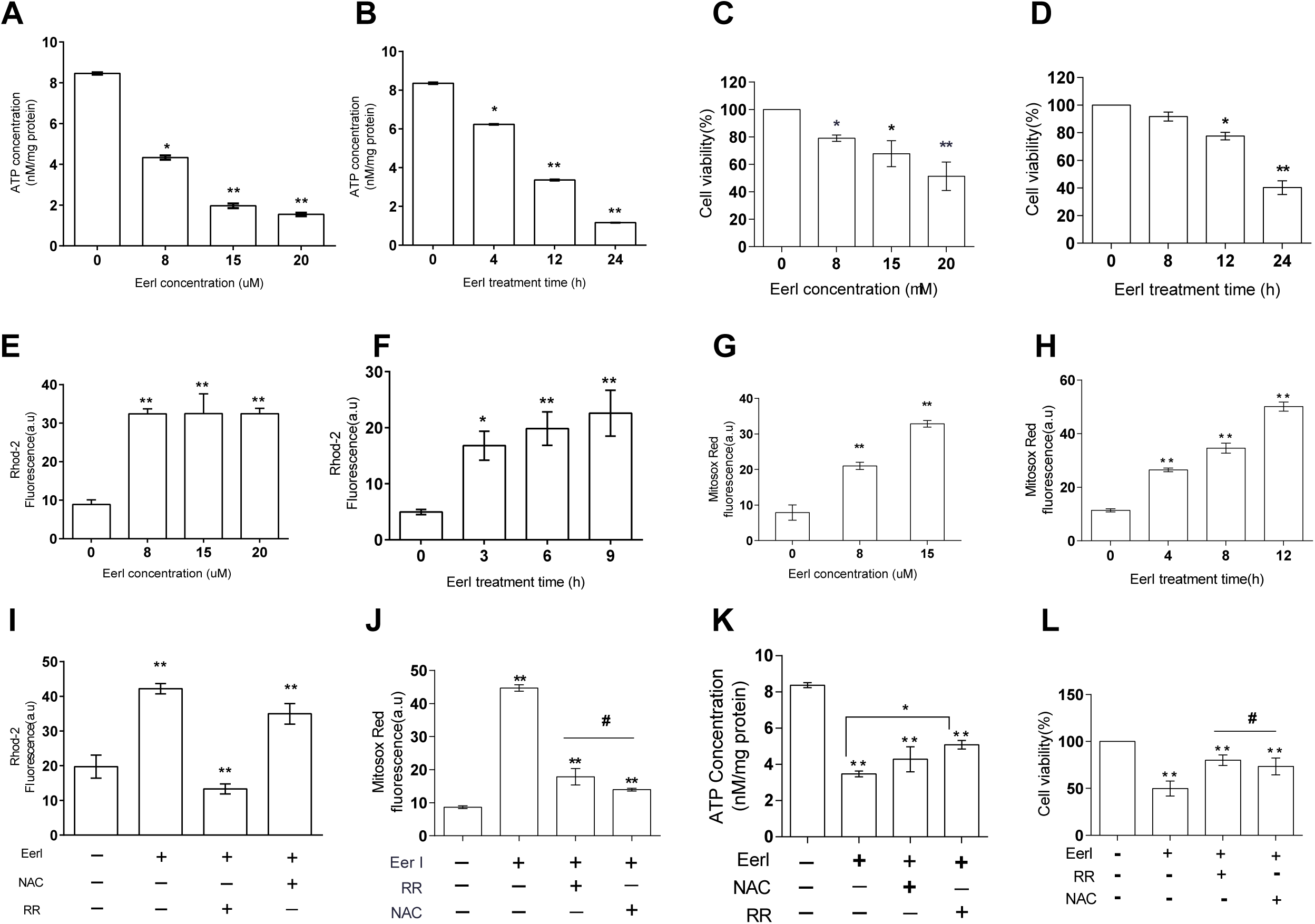
Pharmacological inhibition of ERAD perturbs mitochondrial function and reduces cell viability. HepG2 cells were seeded into 6-well plates at an initial density of 1×10^5^/well and allowed to recover for 12 hours before treating with EerI and other chemical reagents (RR, 2-APB and NAC). Following the treatments, HepG2 cells were analyzed for: (A-B) Cellular ATP concentration; (C-D) Cell viability; (E-F) Mitochondrial Ca2+ and (G-H) Mitochondrial ROS levels. (I-L) Antagonizing effect of RR or NAC on Eer I- induced increase of mitochondrial Ca2+ (I) and ROS (J); and decrease of ATP production (K) and cell viability (L). All data are mean ± SE. *p<0.05; **p<0.01 by Student t-test.

Next we tested the roles of increased of mitochondrial Ca2^+^ and ROS in mediating the effects of EerI on ATP levels and cell viability. HepG2 cells were treated with EerI in absence or presence of the mitochondrial Ca2^+^ uptake inhibitor Ruthenium Red (RR) and the ROS scavenger N-acetylcysteine (NAC). As expected, the stimulatory effects of EerI were reversed by RR in the case of mitochondrial Ca2^+^ and by both RR and NAC in the case of ROS (Fig. 1I-J). More importantly, both RR and NAC attenuated the negative effects of EerI on ATP levels and cell viability (Fig. 1K-L). These results indicate that an effective ERAD is required to maintain normal mitochondrial function and cell viability in mammalian cells.

### Genetic ablation of the core ERAD core protein SEL1L sensitizes HepG2 cells to pro-inflammatory cytokine-induced cell death

Mammalian SEL1L is essential for the formation of the ERAD complex on the ER membrane. To further assess the role of ERAD for normal mitochondrial function, we used CRISPR/Cas9 based gene editing to generate 3 independent SEL1L-deficient HepG2 (Sel1l^-/-^) HepG2 cell lines (Fig 2A-B). While the SEL1L-deficient cell lines were viable and morphologically undistinguishable from the parental HepG2 cell line, they showed markedly higher rate of cell death during long-term culture as determined by real-time monitoring of propidium iodide (PI) staining (Fig. 2C).

**Figure 2.**
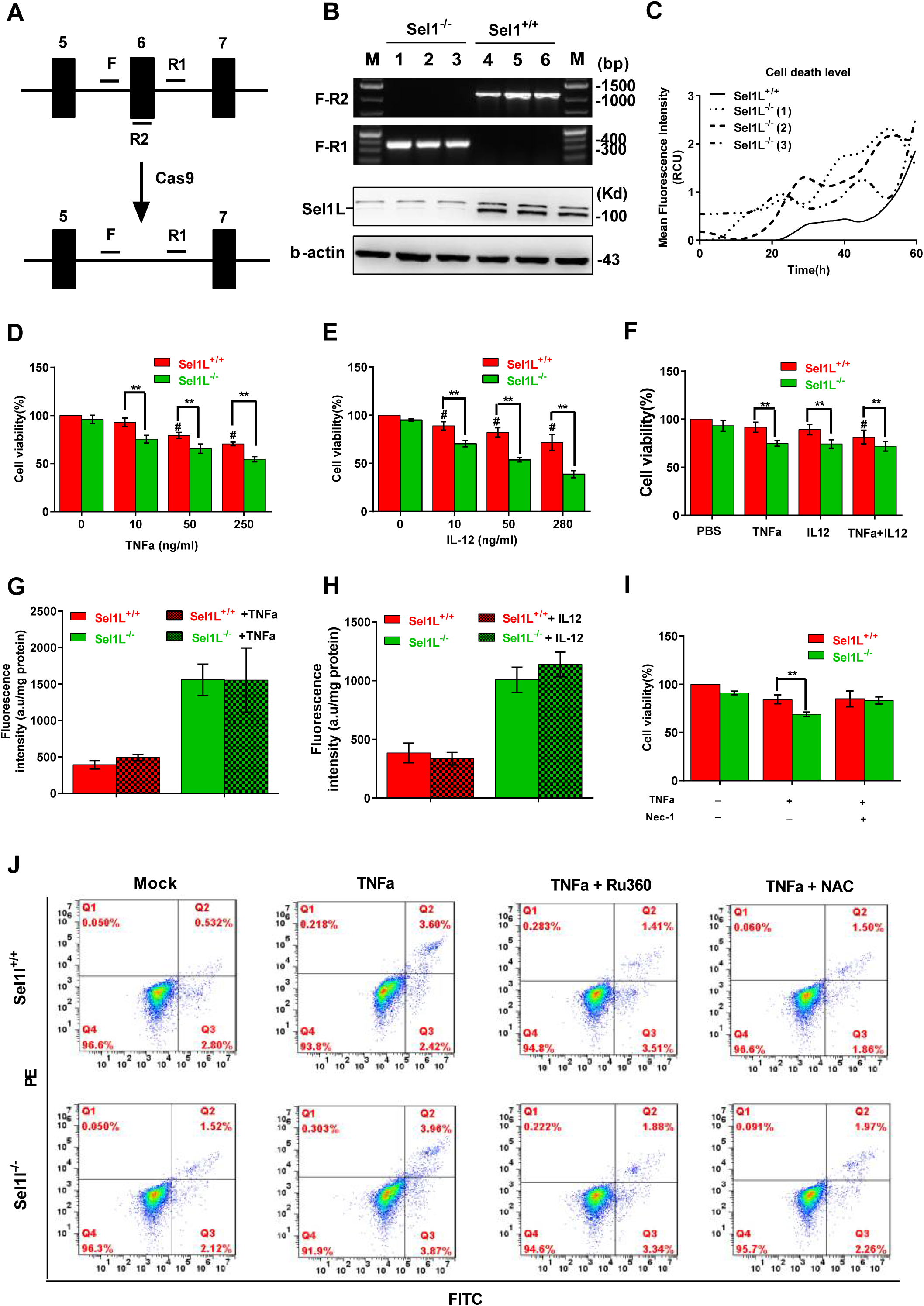
Genetic ablation of ERAD sensitizes HepG2 cells to TNFα and IL-12 induced cell death. (A) Diagram illustrating CRISPR/Cas9-mediated editing of the Sel1L gene in HepG2 cells. F, R1 and R2 represent PCR primers used for genotyping of Sel1L-edited cells. Black bars represent exons. (B) Top: PCR analysis of Sel1L-normal (Sel1L^+/+^) and deleted (Sel1L^-^/-) cells. Bottom: Western blotting analysis of SEL1L expression in Sel1L^+/+^ and Sel1L^-/-^ cells; the upper band is a non-specific band. (C) Average cell death level of Sel1L^+/+^ and Sel1L^-/-^ cells as indicated by the mean PI fluorescence intensity. Sel1L^-/-^ (1-3) represent three independent HepG2 cell clones. (D-F) MTT assays of Sel1L^+/+^ and Sel1L^-/-^ cells after treatment with increased concentrations of TNFα (D), IL-12 (E) and TNFα + IL-12 (F). (G-H) Fluorescence reporter assay of ER stress in TNFα (G) and IL-12 (H) treated Sel1L^+/+^ and Sel1L^-/-^ cells. (I) MTT assay of Sel1L^+/+^ and Sel1L^-/-^ cells after treatment with TNFα or TNFα + Nec-1. (L) FITC and PE dual channel FACS assay of Sel1L^+/+^ and Sel1L^-/-^ cells after treatment with PBS (Mock), TNFα, TNFα + Ru360 or TNFα + NAC. All data are mean ± SE.*p<0.05; **p<0.01 by Student t-test.

A number of pro-inflammatory cytokines promote cell death in diseases affecting the liver and other tissues. To assess whether ERAD deficiency exacerbates cytokine-mediated cell death, WT and SEL1L-deficient HepG2 cells were treated with the pro-inflammatory cytokines TNFα, IL-1α, IL-6 and IL-12) and followed by an assessment of cell viability by the MTT assay. SEL1L-deficient HepG2 cells suffered a higher level of cell death than WT HepG2 cells when incubated with doses as low as 10 ng/ml of either TNFα or IL-12 (Fig 2D and 2E, respectively). Incubation with both TNFa and IL-12 did not yield synergistic effect (Fig 2F). In contrast, no difference in viability was observed between WT and SEL1L-deficient cells incubated with either IL-1α or IL-6 and IL-22 (Fig. S1).

Next we investigated possible mechanisms accounting for the increased susceptibility of SEL1L-deficient cells to the cell death promoting effects of TNFα and IL-12. We first monitored fluorescence in cells transfected with the ER stress reporter plasmid pCAX-F- XBPDBD-venus. As we have previously shown in other cells [15], SEL1L deficiency increased ER stress; more importantly, TNFα and IL-12 did not increase fluorescence in either WT or SEL1L-deficient cells (Fig 2G and 2H, respectively), ruling out a role of ER stress in mediating the effects of TNFα or IL-12. In contrast, incubation with the necrosis inhibitor necrostatin-1 (Nec-1) abolished the increased susceptibility of Sel1l^-/-^ cells to the death-promoting effects of TNFα (Fig 2I). Finally we used annexin V (FITC) and propidium iodide (PE) double dye-staining followed by FACS to ask whether NAC and Ru360 could correct the increased susceptibility of Sel1l^-/-^ cells to TNFα. As shown in Fig 2J, TNFα incubation induced a higher percentage of FITC/PE dual positive cells in Sel1l^-/-^ than in WT population; more importantly, NAC or Ru360 significantly attenuated the death-promoting effects of TNFα. Overall, these results indicate that ERAD-deficient cells are more susceptible to the necrosis-promoting effects of TNFα and a portion of this increased susceptibility is accounted for by increased mitochondrial Ca2+ and ROS levels.

### ERAD deficiency impairs mitochondrial morphology and cellular bioenergetics of HepG2 cells

Mitochondria were imaged using a green fluorescent reporter targeted to the mitochondrial outer membrane [15]. Mitochondria of EerI-treated and SEL1L-deficient cells appeared more rounded than those in control cells (Fig 3B-C vs A). To determine whether these morphological changes were associated with altered function, we stained cells with the lipophilic cationic fluorescent dye JC-1 followed by dual channel FACS analysis. The ratio of high to low (H/L) fluorescence signals was substantially lower in Sel1L-deficient than in WT HepG2 cells indicating a reduction of mitochondrial membrane potential (MMP) in Sel1l^-/-^ cells (Fig 3D).

**Figure 3.**
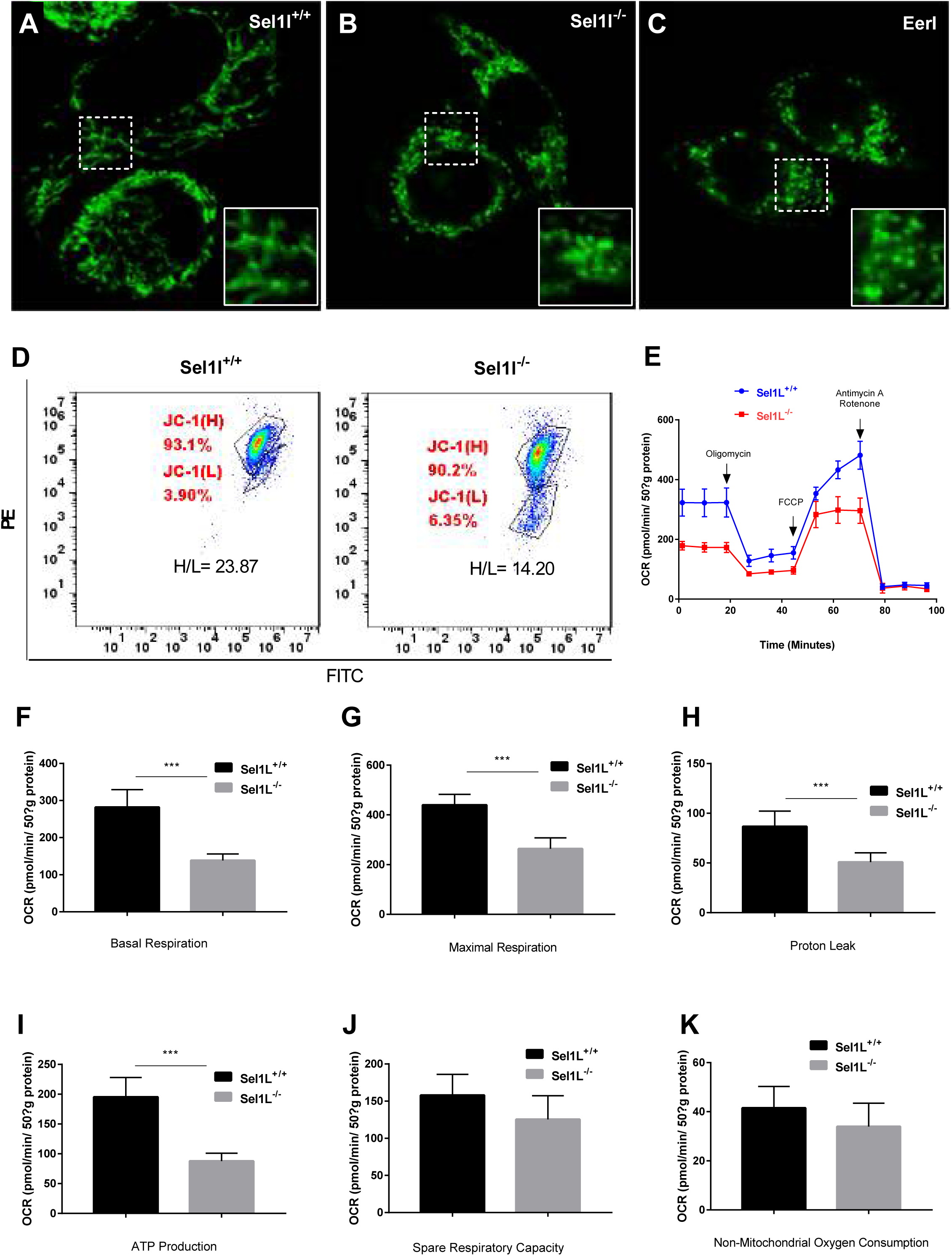
ERAD deficiency alters the morphology and function of mitochondria in HepG2 cells. (A-B) Fluorescent images of Sel1L^+/+^ (A) and Sel1L^-/-^ (B) cells or Eer I-treated Sel1L^+/+^ cells transfected with plasmid expressing a mitochondria-targeted green fluorescent protein (GFP) reporter. Inset at the bottom right of each panel represents a magnified view of the dash line marked area. (D) FITC and PE dual channel FACS analysis of Sel1L^+/+^ and Sel1L^-/-^ cells stained with JC-1. H/L represents mitochondrial membrane potential. (E) Oxygen consumption rate (OCR) profiles (basal or in the presence of oligomycin, FCCP, Antimycin A and Rotenone) of Sel1L^+/+^ and Sel1L^-/-^ cells. (F-K) Quantification of basal (F) and maximal (G) respiration, proton leak (H), ATP production (I), spare respiration capacity (J) and non-mitochondrial oxygen consumption (K) of Sel1L^+/+^ and Sel1L^-/-^ cells. Data are mean ± SE. *p<0.05; **p<0.01 by Student t-test.

We next used Seahorse XF analysis to assess the effects of ERAD deficiency on cellular energetics. Sel1L-deficient cells showed an altered oxygen consumption rate (OCR) profile (Fig. 3E), with significant reductions in basal and maximal respiration rate, proton leak and ATP production significantly reduced (Fig. 3F-I), whereas spare respiratory capacity and non-mitochondrial respiration rate were not altered (Fig 3J-K). In contrast, extracellular acidification rate (ECAR), an index of glycolysis, did not differ between control and Sel1l^-/-^ cells (Fig S2). Overall, these results suggest that ERAD-deficient cells suffer from a reduction of oxidative phosphorylation (OXPHOS), and as a consequence, reduced MMP.

### ERAD deficiency promotes mitochondrial outer membrane permeabilization (MOMP) and sensitizes cells to proinflammatory cytokine induced cell death

Next, we analyzed the effect of ERAD-deficiency and TNFα on the abundance and intracellular distributions of proteins involved in necrosis and apoptosis. Under basal conditions, while there was no difference between control and Sel1L-deficient HepG2 cells in the abundance total c-June N-terminal kinase (JNK), total cytochrome c (Cyto c) and receptor-interacting serine/threonine protein kinase 1(RIPK1), but the latter had higher levels of phosphorylated p54 JNK (p54/p-JNK) and cytoplasmic cyto c (c-Cyto c) (Fig 4A left panels and 4B-C). Upon incubation with TNFα, p54/pJNK and C-cyto c contents in either cell types, but to a greater extent in Sel1L-deficient than in WT cells (Fig 4A middle panels and 4B-C), and this effect was attenuated by conincubation with the JNK inhibitor SP600125 (Fig 4A right panels and 4B-C). Given the increased level of cytochrome c in the cytoplasm, we measured the activity of Caspase 3 and 8. Consistent with the Western blotting result showing increased cytoplasmic content of cyto c, enzymatic analysis showed increased caspase 3 but not caspase 8 activity in Sel1l^-/-^ HepG2 cells, and this pattern was exacerbated by treatment with TNFα (Fig 4D-E). Together, these data indicate that ERAD deficiency promotes mitochondrial outer membrane permeabilization (MOMP) leading to increased release of cytochrome c, and this process is exacerbated in the presence of the proinflammatory cytokine TNFα.

**Figure 4.**
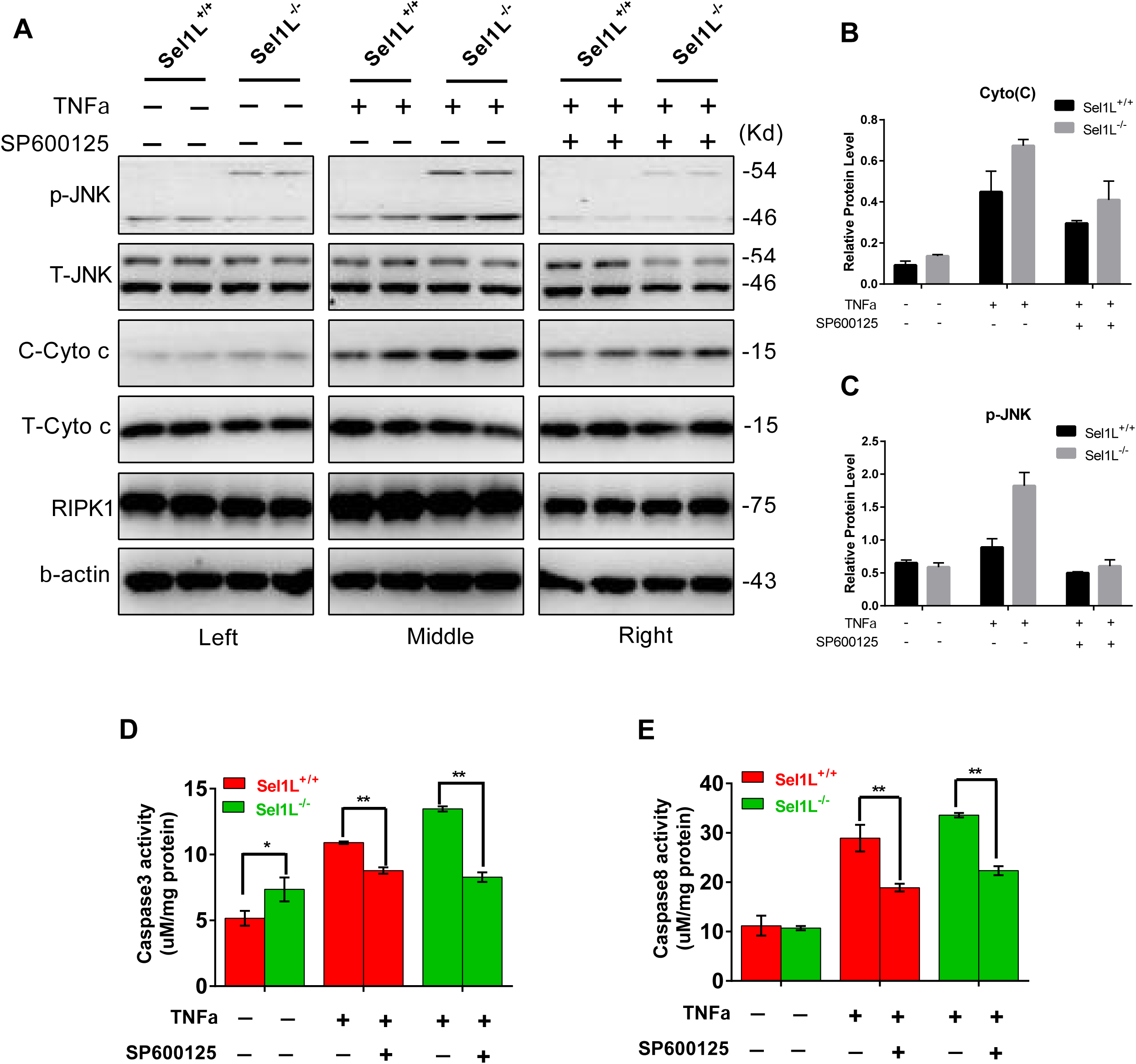
ERAD deficiency induces increased mitochondrial outer membrane permeability and activation of apoptosis. (A) Western blotting analysis of RIPK1, total and cytoplasmic cytochrome c (T- and C-cyto c), total and phosphorylated JNK (T- and p-JNK) in Sel1L^+/+^ and Sel1L^-/-^ cells treated with TNFα or TNFα + SP600125. Β-actin was used as a loading control. (B) Quantification of cytoplasmic cytochrome c and phosphorylated JNK in Sel1L^+/+^ and Sel1L^-/-^ cells following TNFα or TNFα + SP600125 treatment. (D-E) ELISA analysis of Caspase 3 (D) and Caspase 8 (E) activity in TNFα or TNFα + SP600125-treated Sel1L^+/+^ and Sel1L^-/-^ cells. Data are mean ± SE. *p<0.05; **p<0.01 by Student t-test.

### ERAD deficiency causes transcriptional rewiring that disrupts multiple mitochondrial activities but activates several pathways related to ‘stemness’ property of cells

Finally, we used RNAseq on WT and ERAD-deficient cells to obtain a global view of the consequences of ERAD deficiency on gene expression. This analysis identified 1216 differentially expressed genes (DEGs), including 199 up-regulated and 1017 down-regulated genes. By gene enrichment analysis for Gene Ontology biological process and biological pathway, the down-regulated DEGs were found to be significantly enriched in the functional categories including mitochondrial structure and function (mitochondrial ATP synthesis coupled electron transport, mitochondrion organization, mitochondrial respiratory chain complex assembly, mitochondrial translation, and mitochondrial transmembrane transport), endoplasmic reticulum function (protein targeting to ER and establishment of protein localization to endoplasmic reticulum), and metabolic process related to mitochondria or endoplasmic reticulum (oxidative phosphorylation, respiratory electron transport, oxidation−reduction process, and metabolic pathways) (Fig 5A left, 5B). The up-regulated genes were overrepresented in epithelial cell adhesion, P53 signaling, MAPK signaling), mitochondrial structure and function (mitochondrial fragmentation, negative regulation of mitochondrial Ca2+ concentration, regulation of Ca2+ import, Ca2+ channels), endoplasmic reticulum function (protein exit from the ER, protein processing in the ER, unfolded protein response) (Fig 5A right, 5C). Overall, these results indicate that ERAD deficiency caused a global reorganization of mitochondrial activities.

To further understand the pathophysiological significance of the obtained molecular signature in ERAD-deficient HepG2 cells, we performed gene set enrichment analysis (GSEA) by comparing the identified DEGs to priori-established gene modules. As expected, genes of a reported mitochondrial gene module [27] were overrepresented in the down-regulated DEGs in ERAD-deficient HepG2 cells (Fig 5B). Intriguingly, the Jak2 signaling pathway-associated or the so called ‘stemness’ genes [28, 29] were highly represented in the up-regulated DEGs (Fig 5C). Together, these results indicate that ERAD deficiency caused a global reorganization of transcription that disrupts multiple mitochondrial activities and activates several pathways associated with the stemness or invasiveness property of eukaryotic cells.

**Figure 5.**
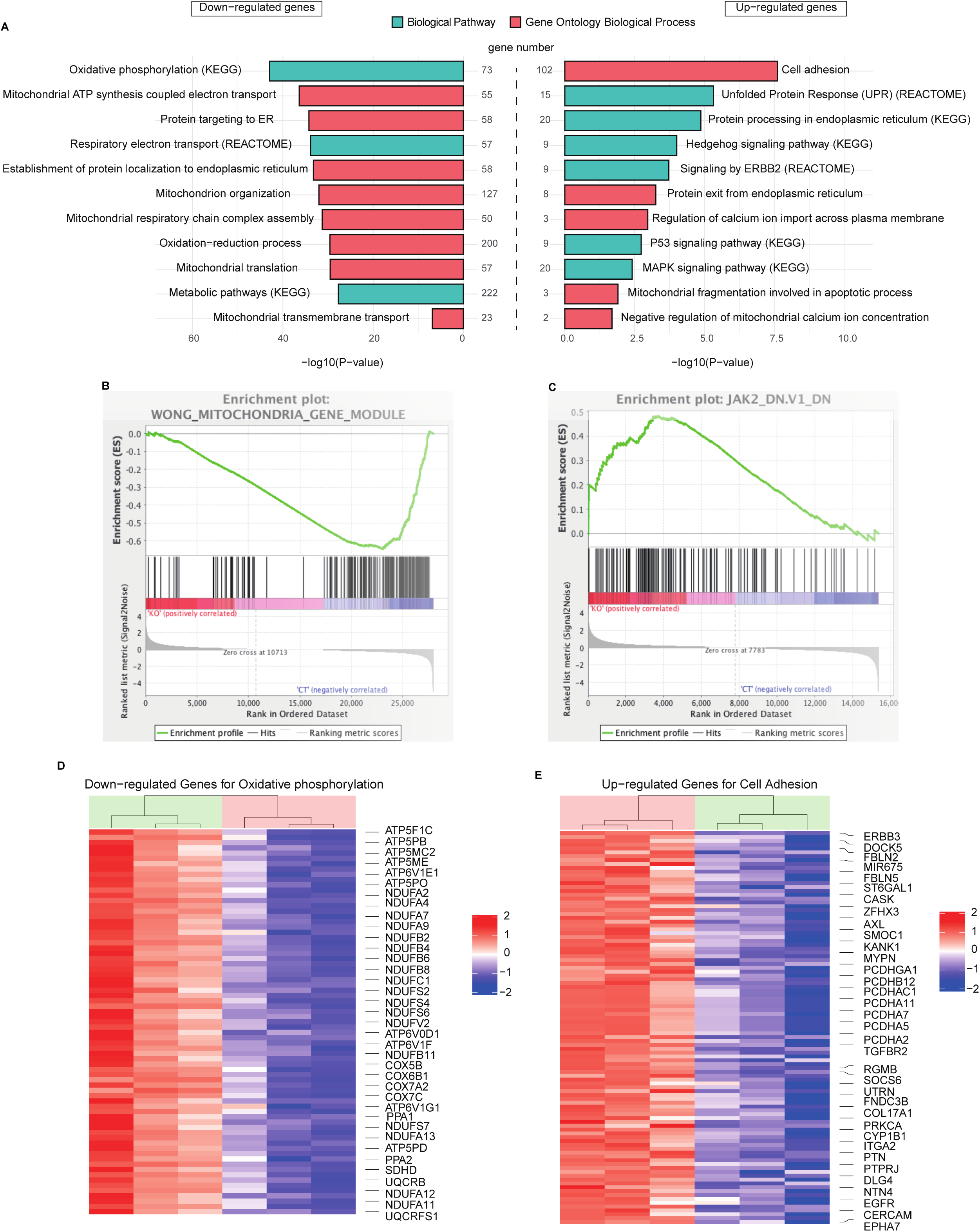
ERAD deficiency causes overhauling of mitochondrial activities and enhances the ‘stemness’ property of HepG2 cells. Sel1L^+/+^ and Sel1L^-/-^ cells were seeded at 1×10^5^/well in 6-well plates and cultured in a humidified CO_2_ incubator for 36 hours before being subjected to total RAN extraction using the TRIzol kit. Three replicates of RNA sample for each genotype were used for RNA sequencing. (A) Gene Ontology biological process (red) and biological pathways enrichment (green) analysis of down-regulated (left) and up-regulated (right) genes in Sel1L^-/-^ cells. (B-C) Representative heatmaps of down-regulated (B) and up-regulated (C) genes in Sel1L^-/-^ cells. Colors indicate scaled expression of individual genes. (D-E) Gene set enrichment plot of down-regulated (D) and up-regulated (E) genes in Sel1L^-/-^ cells. The reference gene modules used in D and E are the Wong mitochondria gene module (https://www.gsea-msigdb.org/gsea/msigdb/cards/WONG_MITOCHONDRIA_GENE_MODULE) and the JAK2 pathway module (https://www.gsea-msigdb.org/gsea/msigdb/cards/JAK2_DN.V1_DN, respectively. Each vertical bar on the x- axis in D and E represents a denoted gene in the specific gene set.

## Discussion

The endoplasmic reticulum and mitochondria are physically and functionally integrated with each other to coordinate an array of metabolic and biosynthetic functions. Stress or dysfunction in one compartment is likely to affect the other and the overall function status of the cell. Here, we provide evidence that deficiency of the ER protein quality control system impacts mitochondrial function in mammalian cells. We show that pharmacological inhibition or genetic disruption of ERAD function decrease the viability of hepatic cell line HepG2 and sensitize them to TNFα-induced cell death. ERAD deficiency induces Ca2+ overflow from the ER to mitochondria, causing over-production of ROS and subsequent damage to the mitochondrial outer membrane. ERAD-deficient cells had decreased ATP production, reduced mitochondrial transmembrane potential and increased outer membrane permeability and, most notably, transcriptome analysis indicates that ERAD deficiency caused a global reorganization of the transcription program. These results highlight an important role for ERAD in maintaining mitochondrial health and in supporting cell survival.

Increasing evidence links ER stress to the development of chronic human diseases, such as type 2 diabetes mellitus, NAFLD, neurodegeneration and cancer [1-3]. However, the mechanisms whereby ER stress leads to the pathogenesis of various human diseases is incompletely understood. ER stress has been shown to activate the unfolded protein response (UPR) responses or the CHOP-dependent cell death program [5, 6]. However, this current UPR model cannot alone fully explain why ER stress preferentially affects metabolically active organs, such as neurons, the endocrine pancreas and liver (Long et.al., unpublished data). In this study, we show that chronic ER stress due to defective ERAD causes structural and functional damages to mitochondria. Moreover, we demonstrate that blocking mitochondria Ca2+ uptake or removing excessive ROS partially rescues cellular defects of ERAD-deficient cells. Given the essential roles of mitochondria in regulating cellular physiology, metabolism and cell death [30], our results suggest that mitochondrial dysfunction may be an under-appreciated driver of many ER stress-induced pathogenesis. These results also strongly support an increasingly recognized notion that normal cellular and organismal health depends on tightly regulated inter-organelle interaction and communication, particularly under stressed conditions, such as viral infection or obesity [31].

Our results provide some mechanistic evidence to explain how ERAD deficiency leads to mitochondrial dysfunction. The ER makes contact with mitochondria through the formation of MAMs, which are temporary structures required for exchange of Ca2+ and other molecules between these two organelles [32]. We showed that, first, elevated levels of Ca2+ and ROS were detected in mitochondria of Eer1-treated and SEL1L-depleted HepG2 cells. Second, blocking mitochondria Ca2+ uptake by treating ERAD-deficient HepG2 cells with Ru360 (a known MCU inhibitor), or removing excessive mitochondrial ROS with NAC (a mitochondria ROS scavenger), partially restores mitochondrial function of ERAD-deficient HepG2 cells. Furthermore, increased cytochrome c, a mitochondrial matrix protein, was detected in the cytoplasmic compartment of ERAD-deficient HepG2 cells. These results suggest a model whereby ERAD deficiency triggers overflow of Ca2+ from the ER to mitochondria, which in turn enhances ROS production and progressive damage to the mitochondrial respiratory system and membrane structure.

Cell death is a common histological feature of many chronic diseases and inflammatory cytokines have been shown to play an important role during this process [33]. For example, the pro-inflammatory cytokine TNFα contributes to the development and progression of inflammation-related diseases, such as non-alcohol steatohepatitis (NASH), type 2 diabetes and cancer, by promoting necrotic cell death [34-37]. Importantly, we show that TNFα induced cell death is increased in the nascence of ERAD and that this effect could be blocked with the necrosis inhibitor Nec-1. These results indicate that ERAD deficiency exacerbates the cell death-promoting effect of TNFα. While further studies are needed to clarify the exact molecular mechanisms underlying the observed synergy between ERAD deficiency and TNFα on cell death, we speculate that the lower intracellular ATP level, the impaired mitochondrial Ca2+ level and increased mitochondrial outer membrane permeability may be among the contributing factors.

ERAD deficiency had profound effects on gene expression in HepG2 cells. RNA sequencing revealed over 1200 DEGs in ERAD-deficient cells with the majority falling in the category of down-regulated genes. Many of those genes are involved in OXPHOS, ATP synthesis, mitochondrial organization and respiratory chain complex assembly (Fig 5A). It is possible that down-regulation of these genes are a consequence of the activation of the mitochondrial unfolded protein response (UPR^mt^) [38]. A major outcome of the activation of UPR^mt^ pathway is suppression of the transcription of mitochondrial function-associated genes, so that the overall mitochondrial activity would be tuned-down to reduce mitochondrial stress [39]. However, many mitochondrial organization and respiratory chain complex assembly are also down- regulated in ERAD-deficient HepG2 cells. A possible explanation for the down-regulation of these genes is that the damage of mitochondria in ERAD-deficient cells has reached a threshold point of “no-return”, so that a mitochondrial functional collapse had occurred.

Intriguingly, most of the 200 up-regulated genes in Sel1L-deficient cells are associated with either cell adhesion or proliferation. Gene set enrichment analysis showed that these genes overlap significantly with a group of so-called “Stemness” genes in the IL6/STAT3 signaling pathway [28, 29]. While further studies are needed to ascertain whether the ERAD-deficient HepG2 cells have reverted to stem-like or more aggressive cells, these results provide evidence that ERAD deficiency may potentiate mammalian cells towards stem or cancer-like cells, perhaps through influencing their mitochondrial function.

In summary, we demonstrate here that ERAD deficiency decrease the viability of HepG2 cells and sensitizes them to TNFα-induced cell death. ERAD deficiency also enhances the stemness or metastasis property of mammalian cells. Mechanistically, ERAD deficiency induces structural and functional damage of mitochondria by impairing the ability of mitochondrial to maintain Ca2+ and ROS homeostasis. Our findings highlight the importance of a regulated ER-mitochondria interaction in supporting normal cellular and organismal physiology, and have important implications in understanding the molecular mechanisms of chronic diseases, such as T2D, NASH, neurodegeneration and cancer.

## Acknowledgements

We thank Yuanyuan Gao, Jiaojiao Sun and Wenyan Ren for technical assistance; Drs. Masayuki Miura, Ying Xu and Han Wang for expression plasmids or other reagents; Dr. Wensheng Zhang for helpful discussions and critical comments on the manuscript.

## Author contribution

X.Q.Y. performed RNAseq and GSEA analyses; Y.B.H. generated Sel1l-deficient HepG-2 cell lines and performed cell viability assay; Q.Q.L. performed the cellular energetics analysis; G.Y.L performed Western blotting analysis. Y.R.B. provided key reagents and helpful discussions; Q.M.L. conceived experiments and wrote the manuscript; all authors reviewed and approved the manuscript. Q.M.L. is the guarantor of this work and, as such, had full access to all of the data in the study and takes responsibility for the integrity of the data and the accuracy of the data analysis

## Figure Legend

**Figure S1.**
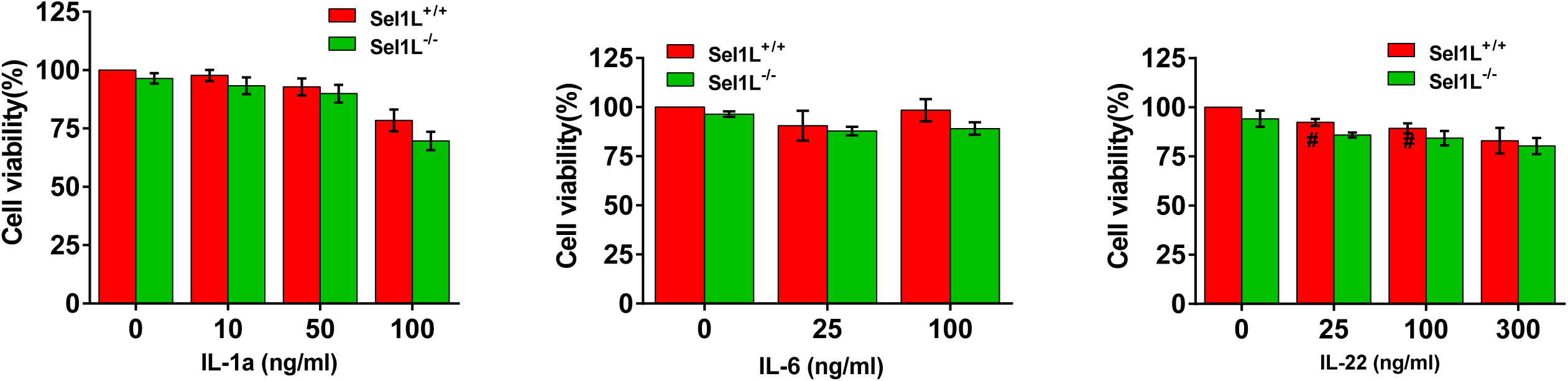
Effects of IL-1α, IL-6 and IL-22 on the viability of Sel1L^+/+^ and Sel1L^-/-^ cells. Sel1L^+/+^ and Sel1L^-/-^ cells were treated for 24 hours with the indicated concentrations of IL-1α, IL-6 and IL-22, and the treated cells were analyzed by MTT assay.

**Figure S2.**
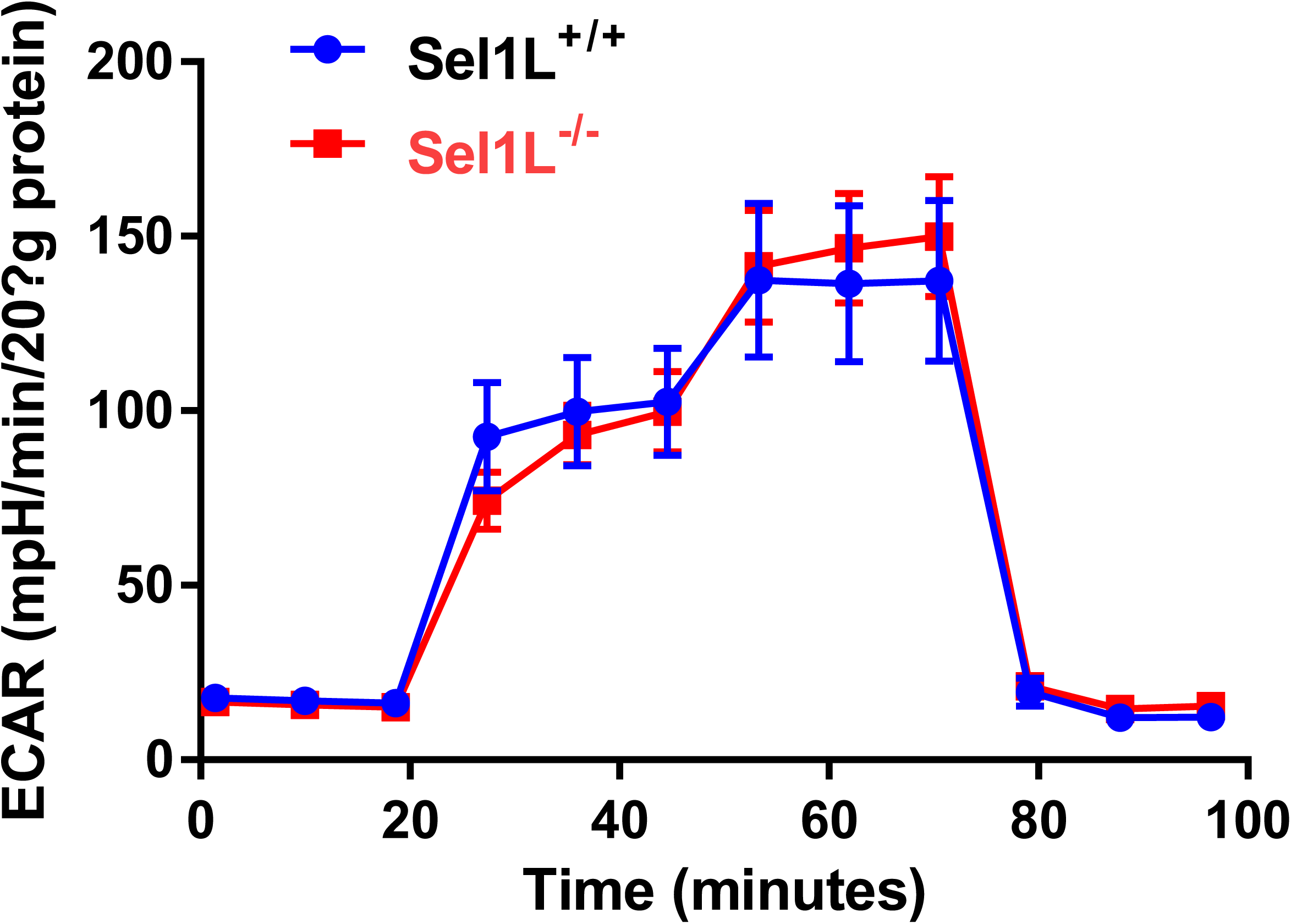
Glycolysis stress test profiles of Sel1L^+/+^ and Sel1L^-/-^ cells. Sel1L^+/+^ and Sel1L^-/-^ cells were seeded at 1×10^4^/well into a 24-well assay plate and allowed to recover for 12 hours. Compounds sequentially injected during the stress test include: glucose, oligomycin and 2-deoxy-glucose (2-DG), which measures glycolysis, glycolytic capacity and allows calculation of glycolytic reserve and nonglycolytic acidification.

